# MS Atlas - A molecular map of brain lesion stages in progressive multiple sclerosis

**DOI:** 10.1101/584920

**Authors:** Tobias Frisch, Maria L. Elkjaer, Richard Reynolds, Tanja Maria Michel, Tim Kacprowski, Mark Burton, Torben A. Kruse, Mads Thomassen, Jan Baumbach, Zsolt Illes

## Abstract

Multiple sclerosis (MS) is a chronic inflammatory neurodegenerative disorder of the central nervous system with an untreatable late progressive phase in a high percentage of patients. Molecular maps of different stages of brain lesion evolution in patients with progressive MS (PMS) are missing but critical for understanding disease development and to identify novel targets to halt progression. We introduce the first MS brain lesion atlas (msatlas.dk), developed to address the current challenges of understanding mechanisms driving the fate of PMS on lesion basis. The MS Atlas gives means for testing research hypotheses, validating candidate biomarkers and drug targets. The MS Atlas data base comprises comprehensive high-quality transcriptomic profiles of 73 brain white matter lesions at different stages of lesion evolution from 10 PMS patients and 25 control white matter samples from five patients with non-neurological disease. The MS Atlas was assembled from next generation RNA sequencing of *post mortem* samples using strict, conservative preprocessing as well as advanced statistical data analysis. It comes with a user-friendly web interface, which allows for querying and interactively analyzing the PMS lesion evolution. It fosters bioinformatics methods for *de novo* network enrichment to extract mechanistic markers for specific lesion types and pathway-based lesion type comparison. We describe examples of how the MS Atlas can be used to extract systems medicine signatures. We also demonstrate how its interface can interactively condense and visualize the atlas’ content. This compendium of mechanistic PMS white matter lesion profiles is an invaluable resource to fuel future multiple sclerosis research and a new basis for treatment development.

## 1 Background

Multiple sclerosis is a chronic inflammatory, demyelinating and neurodegenerative disorder of the central nervous system (CNS) [1]. It is one of the most common causes of neurological disability in young adults [2–4] and the incidence is increasing [5,6]. In about 50% of patients with relapsing multiple sclerosis (RMS), the disease evolves into a progressive phase. At this stage, progression is relentless, and treatments become ineffective. Lesions in the white matter (WM) characterize MS from the early phase. As the disease progresses, quantitative and qualitative changes in the WM can be observed. However, key aspects of PMS pathogenesis are still unsolved, making it challenging to develop treatments. One reason is that direct studies on brain lesions of PMS patients are sparse, and the evolution of acute lesion in the MS brain and their fate are not well characterized. Knowledge from human MS brain lesions is mostly based on candidate gene approaches like immunohistochemistry, microarrays and qPCR. In the last three decades, biobanking of fresh-frozen tissues and advanced technologies in transcriptomics and genomics contributed to more comprehensive studies on the brain material. Unfortunately, most of the generated gene lists do not overlap, which may be due to the use of targeted amplicon sequencing and microarrays, and lack of correction for multiple testing [7–19].

Two recent studies used next generation sequencing [20, 21] on brain tissue but examined only selected lesion types and did not adjust for multiple testing. To allow for identifying systems biology expression signatures that describe brain lesion type formation, evolution and progression, we have assembled the first interactive multiple sclerosis lesion expression map (MS Atlas). At its core sits a database of pre-analyzed whole-genome next-generation RNA sequencing based transcriptomic profiles for stages of lesion formation gathered from 98 *post mortem* human brain samples of 10 patients with PMS and 5 non-neurological disease control cases, including normal appearing WM (NAWM), active, inactive, chronic active and re-myelinating lesions. We applied strict preprocessing and conservative statistics and detected thousands of genes that are significantly differentially expressed during lesion evolution compared to control samples (Figure 1). Our MS Atlas features an online web-based data analysis platform to identify and extract mechanistic pathways and gene sets that distinguish lesion types and are candidate drivers of different lesion type formation.

**Figure 1.**
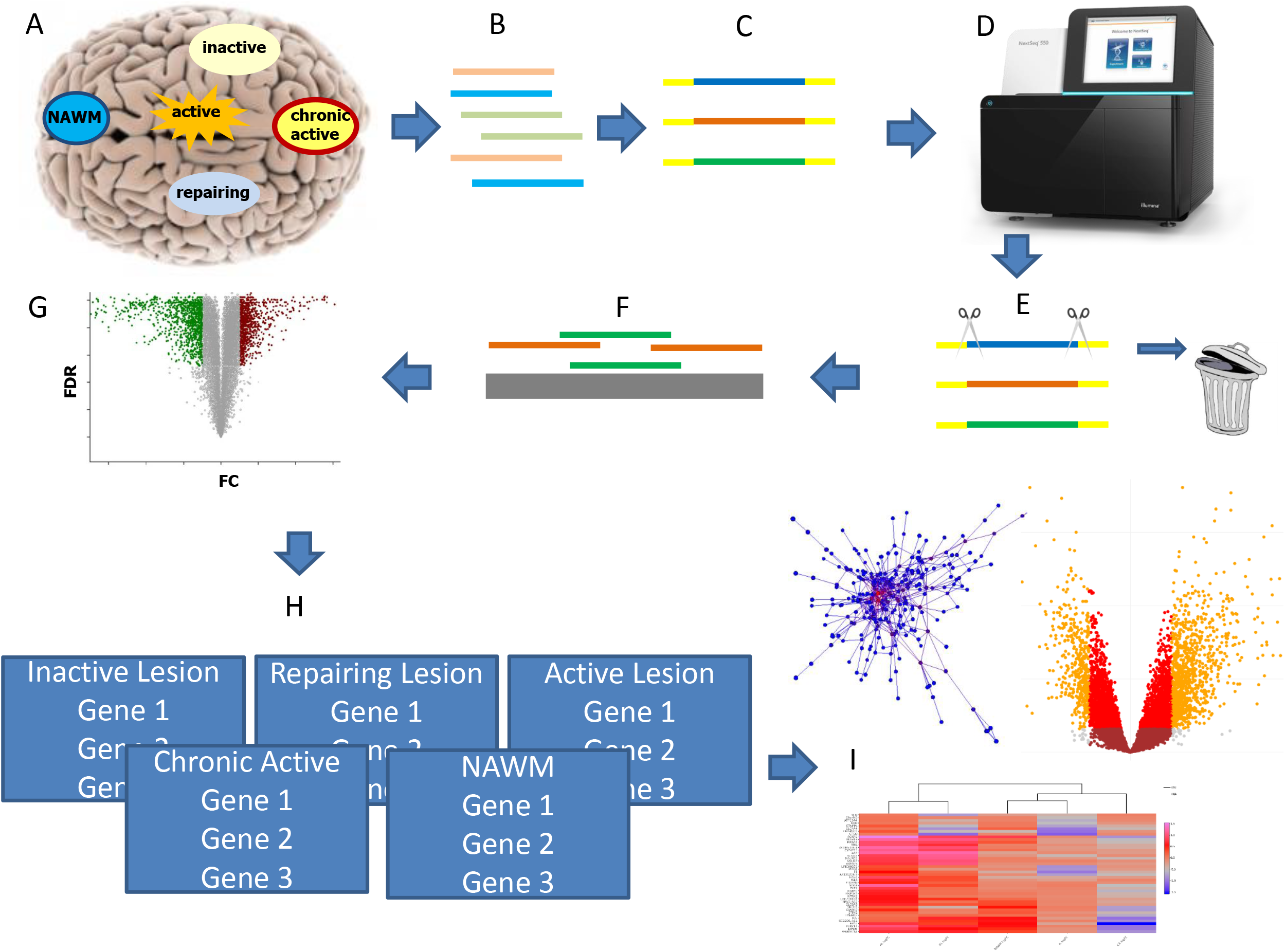
Graphical summary of the study. (A) With immunohistochemistry, we classified different lesion types (normal appearing white matter (NAWM), active, inactive, chronic active, remyelinating), and microdissected 98 of these brain areas from ten progressive multiple sclerosis and five non-neurological disease brains. (B) Total RNA (microRNA, lncRNA, and protein-coding) was extracted and (C) library was generated. (D) The templates were sequenced on a NextSeq550 using paired end, followed by (E) quality control, where all reads below Q20 were removed. (F) Remaining reads were aligned to the human genome and counted. (G) Statistical calculations were performed in R and the significant threshold was set to FDR < 0.05. (H) MS Atlas was implemented based on RShiny offering three major visualizations (I): heatmaps, mechanistic candidate networks, volcano plots.

## 2 Construction and content

### 2.1 Human Postmortem Brain Tissue

Seventy-five snap-frozen tissue blocks from ten PMS patients and 25 blocks from five donors without neurological disease have been obtained from the United Kingdom Multiple Sclerosis Tissue Bank (UKMSTB) at Imperial College London. All tissues were obtained within 30 hours after death. The age of patients at death was 52.4±10.2 years, and the age of the controls was 56.4±14.1 years. We examined 4-10 brain areas/lesions from each brain: altogether 20 normal-appearing white matter areas (7 patients), 17 active lesions (8 patients), 14 inactive lesions (5 patients), 6 remyelinating lesions (4 patients), 17 chronic active lesions (7 patients), and 25 control white matter areas (5 controls).

### 2.2 Immunohistochemistry and lesion classification

Snap-frozen tissue has been sectioned and stained for classification of NAWM, active, inactive and remyelinating lesions based on antibodies against myelin oligodendrocyte glycoprotein (MOG) to detect myelin integrity and human leukocyte antigen D related (HLA-DR+) to characterize the inflammatory state [22]. For staining with VLA4/integrin *α*-4 antibody (ab77528, abcam), we used tissue from one MS patient and control.

### 2.3 RNA extraction from specific histological brain areas

The brain fields of interest were microdissected in a cryostat (10-100 mg/sample). Total RNA has been isolated with miRNeasy Mini Kit from Qiagen, and DNAse I treatment (RNAse-Free DNAse Set, Qiagen) was applied to eliminate genomic DNA interference. RNA concentration and purity have been measured on a NanoDrop spectrophotometer ND-1000 (Thermo Scientific) and the integrity of RNA (RIN) was measured using the Bioanalyzer 2100 (Agilent Technologies). The fragmentation time and cleanup steps during library preparation have been adapted for each sample based on the RIN value.

### 2.4 RNA-seq

One *μ*g of RNA per sample was processed to remove ribosomal RNA followed by library preparation using TruSeq Stranded Total RNA Library Prep Kit with Ribo-Zero Human/Mouse/Rat Set (Illumina). The quality and fragmentation size of the libraries were estimated by High Sensitivity DNA chip on the Agilent 2100 Bioanalyzer and the concentration determined with Qubit dsDNA HS Assay (Life Technologies, Carlsbad, CA). Two pM pooled indexed libraries were loaded into flow cell followed by cluster generation and indexed paired-end sequencing (80+7+80 bp) on Illumina NextSeq500/550 (High Output v2 kit (150 cycles)).

### 2.5 Raw data analysis and quality control

The data were de-multiplexed by the Illumina machine and exported in the FASTQ file format. Afterwards, the read quality for the 100 samples was accessed via FastQC [23]. Trimmomatic [24] was used in order to trim the reads and remove any hypothetical adapter contamination. The software was provided with the correct Illumina adapter sequences and the quality cutoff for the leading/trailing bases as well as for the sliding windows was set to 20. The minimal length of the trimmed reads was set to 17 in order to include potentially present microRNA. In the next step, STAR aligner was utilized for read mapping against the human genome (hg38, downloaded 08.05.2017). The mapped reads were further processed using htseq-count [25] in strict mode in order to access raw read counts for every gene.

Two samples have been excluded during quality control. Some of the remaining 98 samples had to be re-sequenced due to low read count. In total, we ended up with 73 cases and 25 controls.

### 2.6 Statistics

The unique Ensemble IDs were mapped to gene symbols using the R package “org.Hs.eg.db” [26]. In case the package was missing a gene symbol the Ensemble ID was replaced with a unique number. During the mapping process, about 25% of the Ensemble IDs could not be mapped to a gene symbol. Note that the gene symbols are solely used for result visualization while all analyzes have been performed based on the unambiguous Ensemble IDs. We used EdgeR [27] to process the raw read counts and scan for significant genes for the different lesion types. All samples have been normalized for library sizes. Five generalized linear models were trained in order to reveal differentially expressed genes between control (WM) and each of the five lesions types (NAWM, active, chronic active, inactive, remyelinating). All models were adjusted for age and sex. Our models additionally account for lesion distribution, since from every patient multiple samples of the same lesion types have been extracted and used. We obtain, for every lesion, a list of genes with the corresponding log_2_-fold-changes (logFC) and the p-value corrected for multiple testing using FDR-correction (Benjamini-Hochberg) [28].

### 2.7 The MS Atlas database and online analysis platform

The processed data was then integrated into a database, that we make publicly available to the research community together with a web-interface based on RShiny [29]. Besides data download, the platform offers three major visualization tools to compare the different lesion types and extract markers at different levels in the system biology value chain. Heatmaps and volcano plots allow for the extraction of gene panels associated with lesion type. Network enrichment methodology (i.e. Key-PatwhayMiner [30,31]) enables the identification of mechanistic (i.e. sub-network-based) markers. We integrated the human protein-protein interaction network from the IID database [32] filtered for only brain tissue specific interactions with experimental evidence, orthologous mice genes and computational prediction. The MS Atlas online platform produces visualizations on-the-fly using a variety of R packages (Table 1).

**Table 1.** R-packages

| Name | Version |
| --- | --- |
| conflicted [41] | 0.1.0 |
| shiny [42] | 1.1.0 |
| ggplot2 [43] | 2.2.1 |
| plotly [44] | 4.7.1 |
| shinycssloaders [45] | 0.2.0 |
| plyr [46] | 1.8.0 |
| heatmaply [47] | 0.15.0 |
| threejs [48] | 0.3.1 |

## 3 Utility and Discussion

The MS atlas is the first platform for mapping MS lesion expression profiles and the generation and testing of specific hypotheses. In the following, we will introduce the workflow of the online platform and a use-case for its biomedical application.

### 3.1 Workflow

MS Atlas web interface offers three major visualization themes: heatmaps, mechanistic candidate networks and volcano plots. The user can adjust several parameters (Figure 2 A) to choose statistical significance levels of the profiled genes. Initially, the user selects a (set of) lesion type(s) of interest. The user is further asked to choose whether the interest is in up-, down- or overall deregulated genes. To study the evolution and development of lesion types, one might, e.g., filter for genes down-regulated in chronic but upregulated in inactive lesions. In the next, step those genes can be sorted by FDR-corrected p-value and filtered for a minimal logFC value. The user may also directly search for a specific gene of interest and check for its expression changes across different lesion types. The MS Atlas platform will then visualize the gene’s expression across the selected lesions in a heatmap (Figure 2 B). Genes and lesions are ordered based on a hierarchical clustering using the Euclidean distance metric. The color represents logFC (lesion vs. control). The individual logFC of a gene in a lesion is shown in a tool-tip when the mouse is hovering over the corresponding field. All data shown in the heatmap can be exported as CSV file or PNG image.

**Figure 2.**
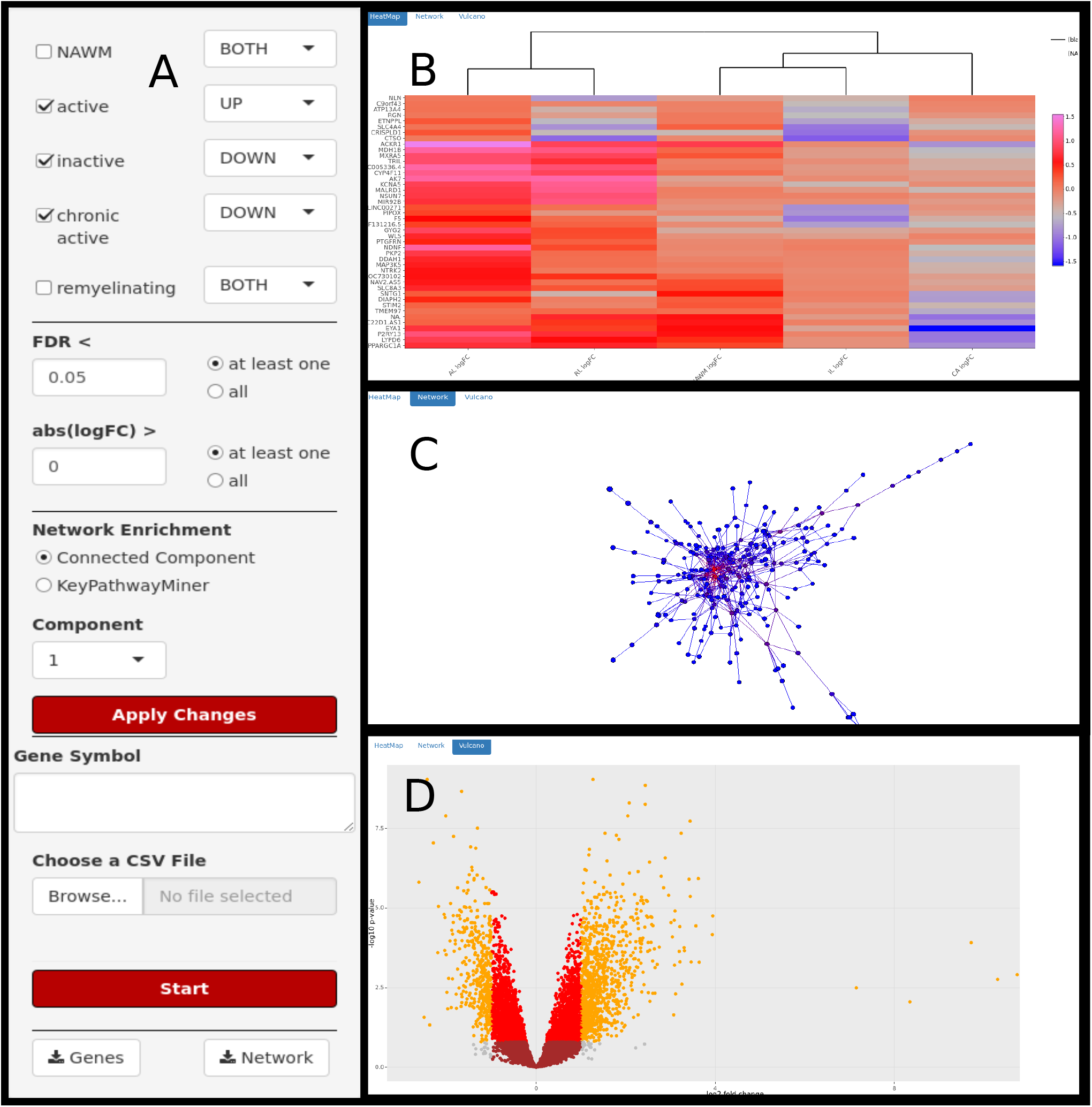
Web-interface of The MS Atlas. (A) The MS Atlas web interface offers adjustment for statistical parameters (e.g. lesion types, FDR, logFC) and subnetwork extraction. Significant genes will be visualized in a (B) heatmap, (C) mechanistic candidate network and (D) volcano plot.

Furthermore, to suggest potential mechanistic markers putatively driving lesion type evolution, selected genes can be projected onto the human protein-protein interaction network (Figure 2 C). Since the network contains over 400,000 interactions and 13,000 genes, displaying the full network would not help extracting useful information. Instead, we integrated the *de novo* network enrichment method KeyPathwayMiner, which extracts sub-networks that distinguish, on a mechanistic level, between MS lesion types and, thus, provides first hints on how lesion evolution is driven and controlled on a systems biology level. We allow to specify a number of exception genes (k), which do not necessarily have to be significantly differentially expressed between lesion types (i.e. outliers) but still play a central role in the interaction network. A mouse klick on a node/gene in the network reveals additional information. The key networks can be exported in SIF format for downstream analyses in Cytoscape [33] or as PNG image file.

Finally, the platform allows for on-the-fly visualization of volcano plots for all genes of a selected lesion (see Figure 2 D), where two thresholds have been chosen (FDR < 0.05 and logFC < 1.5) to color-code the genes accordingly.

### 3.2 Drug target expression during lesion genesis

Natalizumab is a monoclonal antibody used in the treatment of RMS patients [34]. It blocks the alpha4 integrin(VLA-4)-mediated trafficking of pathogenic lymphocytes through the blood-brain barrier, and prevents inflammation in the CNS [35]. While natalizumab is one of the most effective treatments in RMS patients [36], its efficacy in PMS is limited [37]. Our database and validation by immunohistochemistry indicate that VLA-4 is the most highly expressed in active lesions even in the PMS phase, but it is signficantly upregulated in all lesion types compared to the normal-appearing white matter (NAWM) (Figure 3); the limited efficacy in the ASCEND clinical trial of PMS may be related to the increasing number of chronic active lesions in this phase of the disease with less expression of VLA-4 compared to the active lesions [38,39]. Alternatively, additional mechanisms of inflammation that are unrelated to VLA-4 and/or increasing dominance of pathways independent of inflammation may contribute [40].

**Figure 3.**
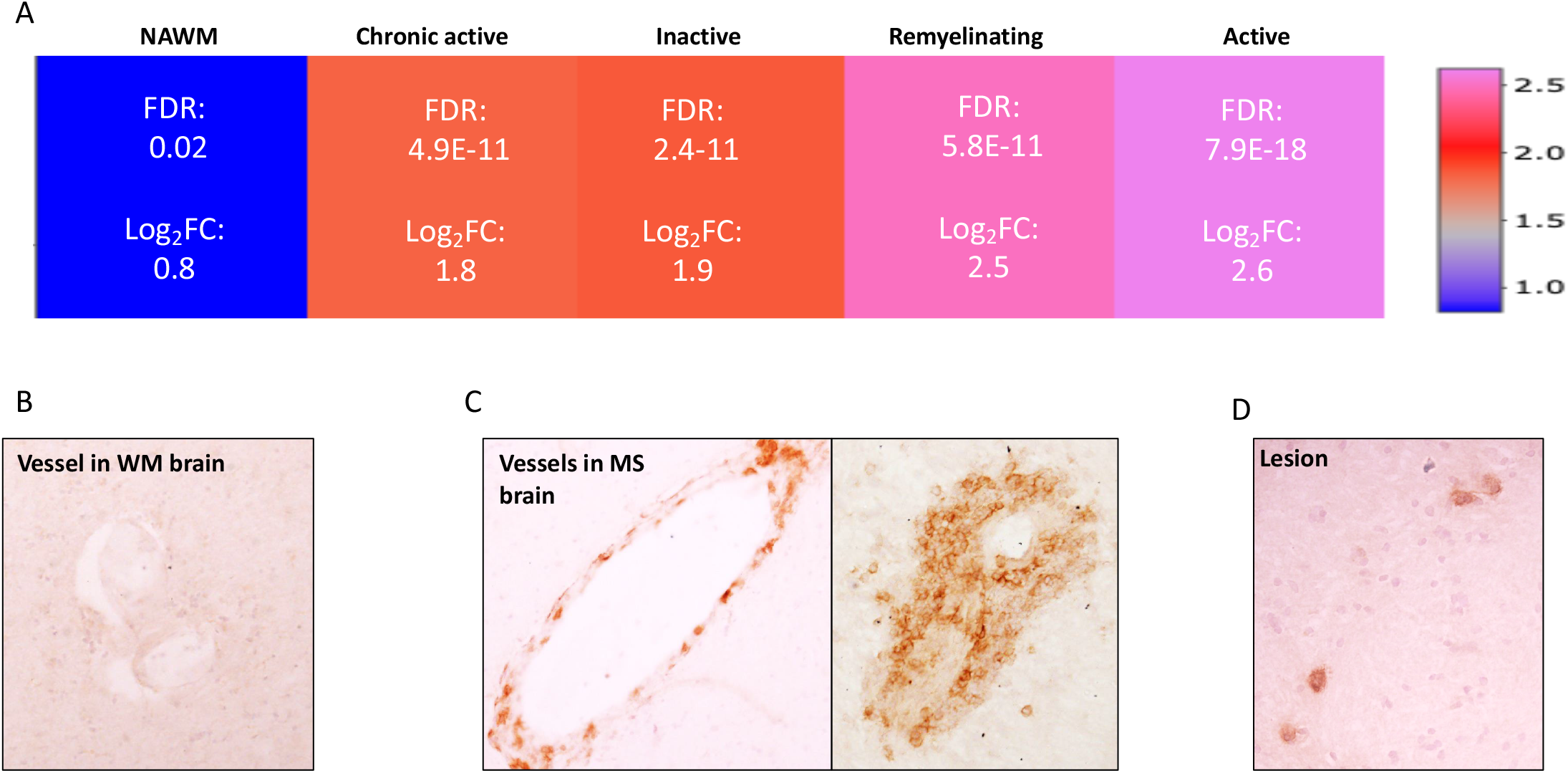
Expression of drug target in the MS Atlas and evaluation on protein level. (A) Gene expression pattern of VLA4 receptor (drug target) in normal appearing white matter (NAWM), active-, inactive-, remyelinating- and chronic active lesion extracted from the MS Atlas. (B) Immunohistochemistry of VLA4 receptor in WM brain tissue, from non-neurological disease (C) High protein expression of VLA4 receptor in vessels of MS brain tissue (D) Detection of VLA4 receptor in parenchyma of SPMS brain tissue

## 4 Conclusion and future development

We present the first atlas of multiple sclerosis transcriptomic brain lesion maps. It is based on total RNA next-generation sequencing of different lesion types in PMS patients compared to WM of non-neurological disease controls. The MS Atlas will fuel future research projects and significantly aid in advancing not only the MS field but also for research in other neurological diseases, as it allows researchers to (i) search for certain molecules of interest as drug targets or biomarkers of brain lesion genesis; (ii) compare gene panels extracted from functional cell or animal studies; (iii) discover mechanistic markers using *de novo* network enrichment from genes of interest in different lesion types. We showed that the drug target known to be only effective in early disease stages is present in the active-but less in the chronic active lesion types characteristic of progressive MS. The MS Atlas is extendible and will continuously be updated with future transcriptome profiles. We also aim to integrate genome-wide methylome data from the same tissues, making it possible for the user to correlate the gene expression with methylation status, and to mine its mechanistic joint effects on lesion evolution.

